# Dynamic inventory of moths of Savitribai Phule Pune University (Pune, India) through crowdsourcing via *iNaturalist*

**DOI:** 10.1101/2021.08.01.454690

**Authors:** Bhalchandra Pujari

## Abstract

We present here the checklist of moths (Lepidoptera: Heterocera) for the campus of Savitribai Phule Pune University, situated in the metropolis of Pune in the state Maharashra in India. We report identification of 189 unique genera along with 154 unique species. Despite the relative small size of the observation area and the location being at the heart of a busy metropolis, the moths were found to be of diverse variety, with 26 different families and 76 tribes.

The identifications of the species was crowd-sourced via iNaturalist.org. An automated program was developed to fetch the identification and generate the list. The program is also being made available open source. As a result the checklist remains relevant all the time with newer identification, correcting existing identification (if required) and taxonomic updates of the future.

## Introduction

The bustling metropolis of Pune is located at the western part of India at 18°31’13”N and 73°51’24”E (Fig 1(a)). The metropolitan area of the city has estimated population of 7.4 million (Wikipedia (2021)) and is home to several education institutes including Savitribai Phule Pune University (formerly known as University of Pune). The city is located at the foothills of Western Ghats (Sahyadri mountain ranges) and is rich in biodiversity.

**Fig 1:**
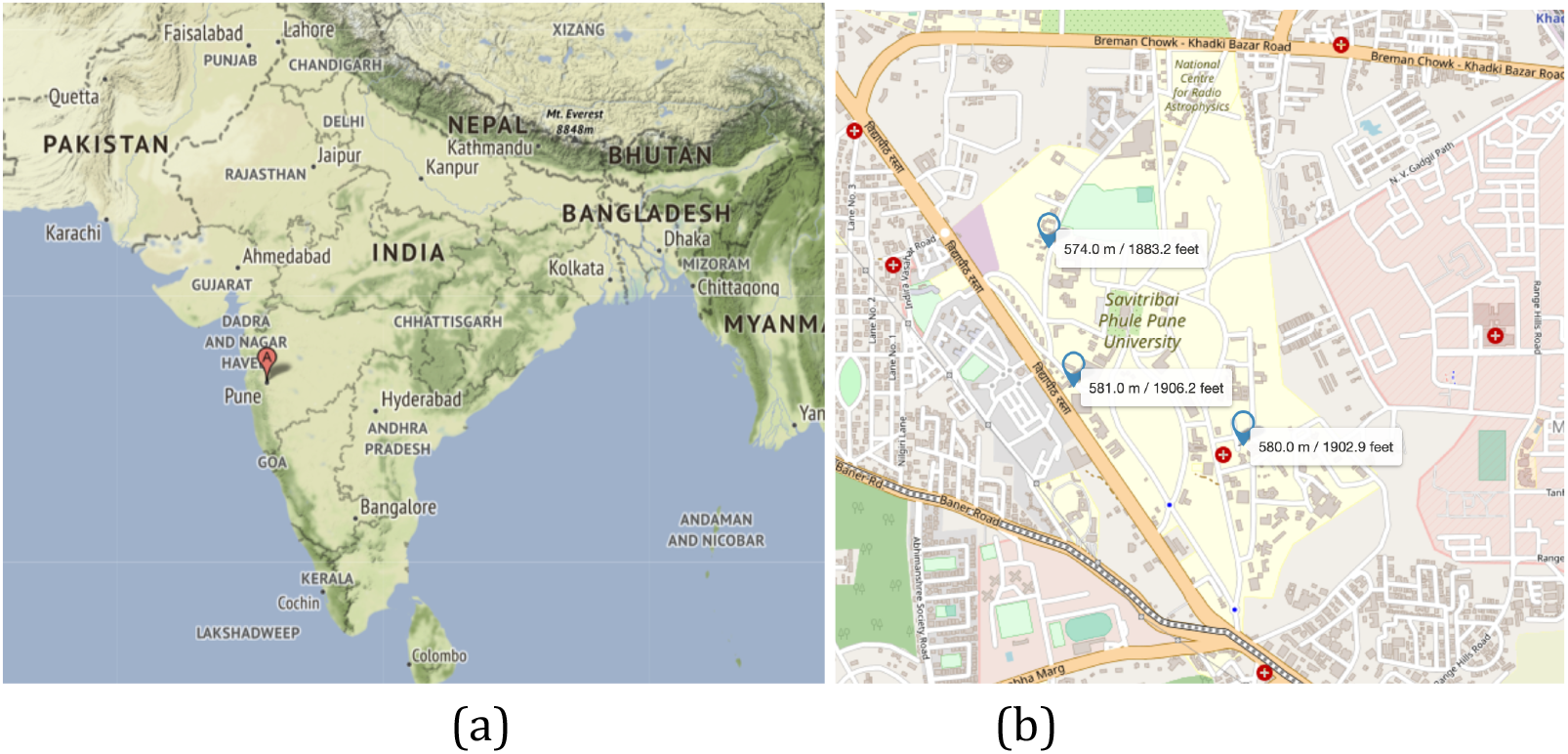
(a) Pune city, shown with red marker, is located at the western part of India in the state of Maharashtra. It lies west of the Western Ghats. (b) Map of the university (shaded in yellow) along with the locations (in blue) where most of the observations were carried out. Courtesy: ACME mapper/Openstreetmap.

Established in 1949 the university is spread across the area of 411 acres and is considered to be a prime institute in India. The campus is located roughly at the elevation of 580 m from mean sea level. Ecologically one the unique features of the university campus is the presence of large numbers of Dalbergia melanoxylon (African blackwood) trees, believed to have been introduced in colonial times by British government. Such a large population of these trees is not reported anywhere in India. Apart from Dalbergia melanoxylon there are a handful of introduced species like Couroupita guianensis (cannon-ball), guazuma ulmifolia (West Indian elm), Libidibia coriaria (divi-divi), adansonia (baobabs) among others. The campus also hosts multiple native trees mainly ficus benghalensis (Indian banyan), ficus religiosa (sacred fig), Cassia fistula (golden rain), jacaranda mimosifolia (blue jacaranda) etc.

Despite the rich biodiversity we could not find the comprehensive checklist for the campus or even broader metropolis region. One of the most important works in the nearby region on lepidoptera inventory is due to Shubhalaxmi et. al. (2011) wherein the authors have explored the said fauna in Western Ghats. Their areas of observations has been mainly the hilly region with dense forest cover. They have reported a rich fauna but it is expected that an urban environment is unlikely to support same diversity. The other records do exist from northern Maharashtra (Gurule and Nikam (2011, 2013), but the terrain and the climate is significantly differ from that in Pune. Therefore it was imperative to document the Lepidoptera fauna in its current state for the university campus.

## Methodology

To generate the checklist we decided to utilise the power of crowd sourcing using modern technologies. With its powerful Artificial Intelligence engine as well as presence of various taxonomists, iNaturalist (iNaturalist(2021)) was our choice of technology. iNaturalist is a platform for scientists and citizens alike to share the observations from all across the globe. The records are available to public and are naturally peer-reviewed at all the times. Additionally the records from iNaturalist is also shared with Global Biodiversity Information Facility (GBIF(2021)) for broader scientific use.

For data collections we carried more than two years of observations from 2019 to 2021 and ongoing. We periodically observed the walls lit with 60W compact fluorescent lamps located mainly at three locations on the campus: 1) near Jaykar library, 2) Faculty house and 3) department of scientific computing modeling & simulation. Some insignificant number of observations were also made at other location across the campus. To be the non-invasive, no separate traps were set up; neither any moth was captured for the observation. As a result some of the identification had to be limited to genera level.

## Results

Before we start discussing the findings from the observation it is worth pointing out that results we are about to present are dynamic in nature. That means over the period of time some updates are expected due to addition of new observation, correction of existing identification or taxonomic changes of future. This ensures it will be continually relevant and never be outdated. In order to stay up-to-date the dynamic version of the checklist is also made available on https://gitlab.com/bspujari/sppu_moths. Additionally an interactive webpage is also maintained at [author’s webpge](http://cms.unipune.ac.in/~bspujari/Pune_university_Lepidoptera/)^2^. Moreover for sake of accessibility the complete Python program that fetches the data from iNaturalist and compiles the list, is made available - open source^3^ - on both of these pages.

At the time of writing of this manuscript, total of 880 observations were made with 737 identification at various level of taxonomy. The list of uniquely identified species is collected in Table 1, at the end of this section. The table also show an “Obs ID” with each observations which is a unique identification number assigned by iNaturalist. One can simply use inaturalist.org/observations/<<Obs ID>> to reach the dedicated observation page which includes the details of taxa as well as media associated with the observation. Additionally any taxonomic updates are also reflected on the page, for example, <u>inaturalist.org/observations/30996315</u> shows how Pyrausta panopealis was swapped to Pyrausta phoenicealis, with the reasoning given on inaturalist.org/taxon_changes/64986.

**Table 1:**
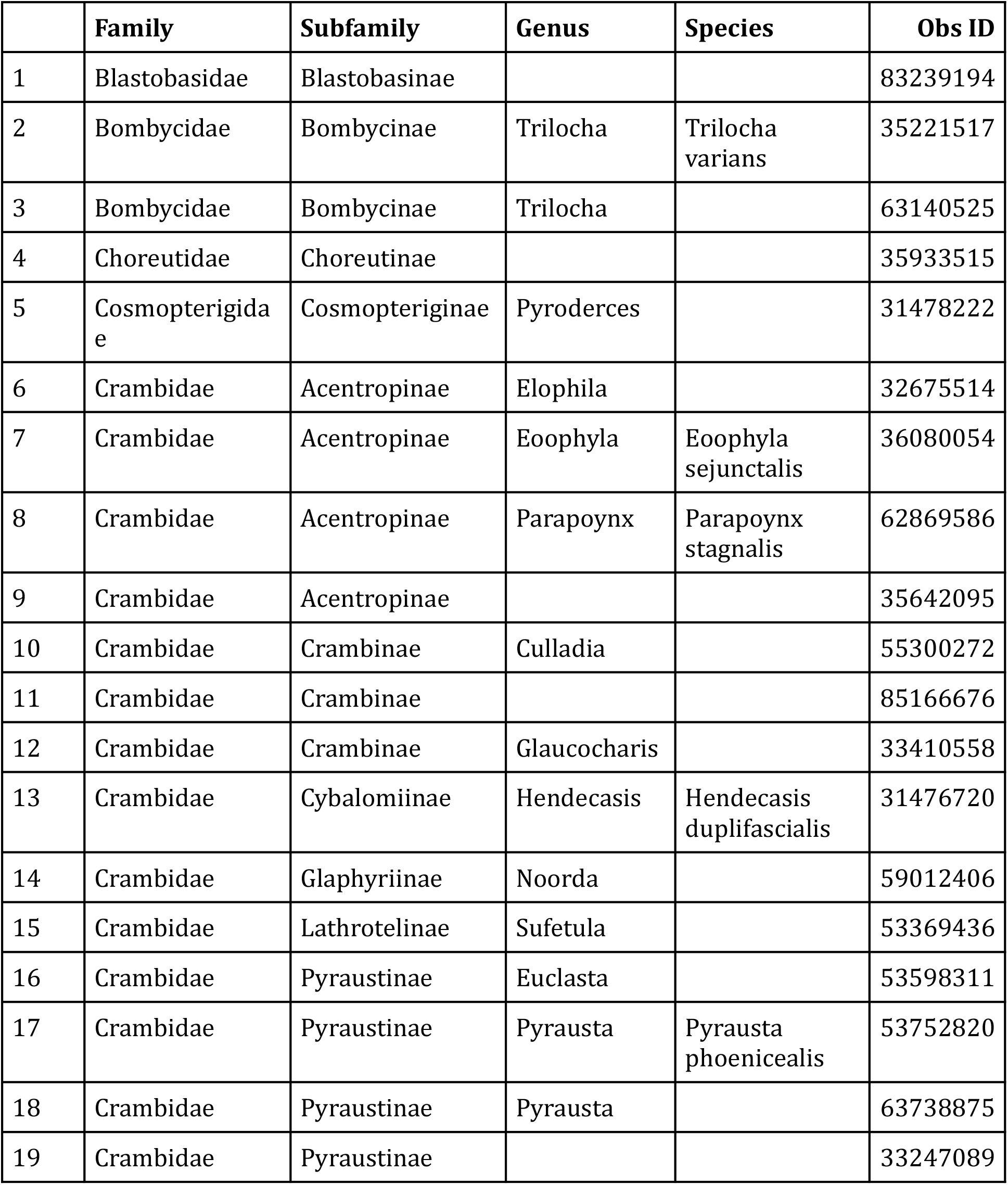
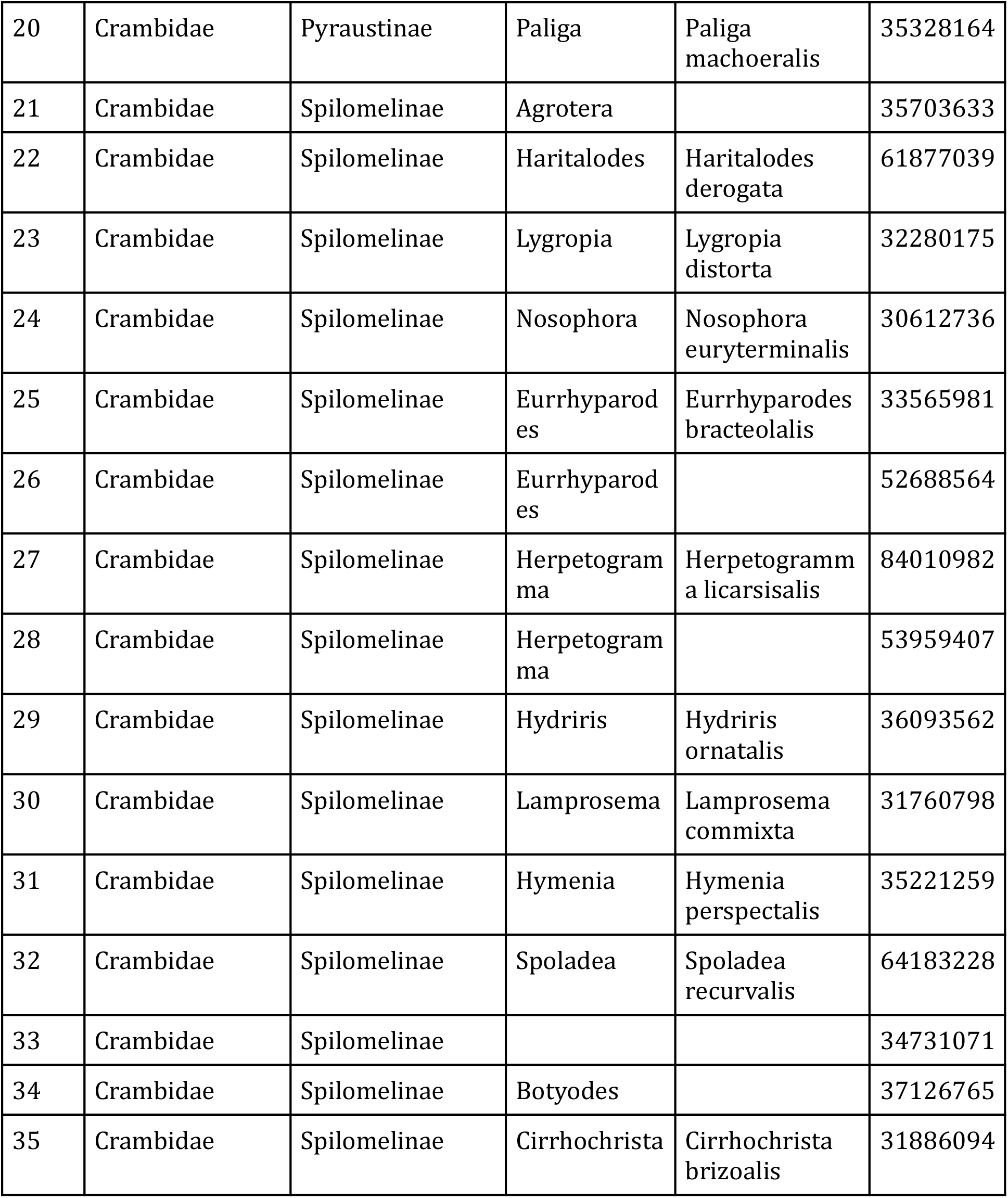
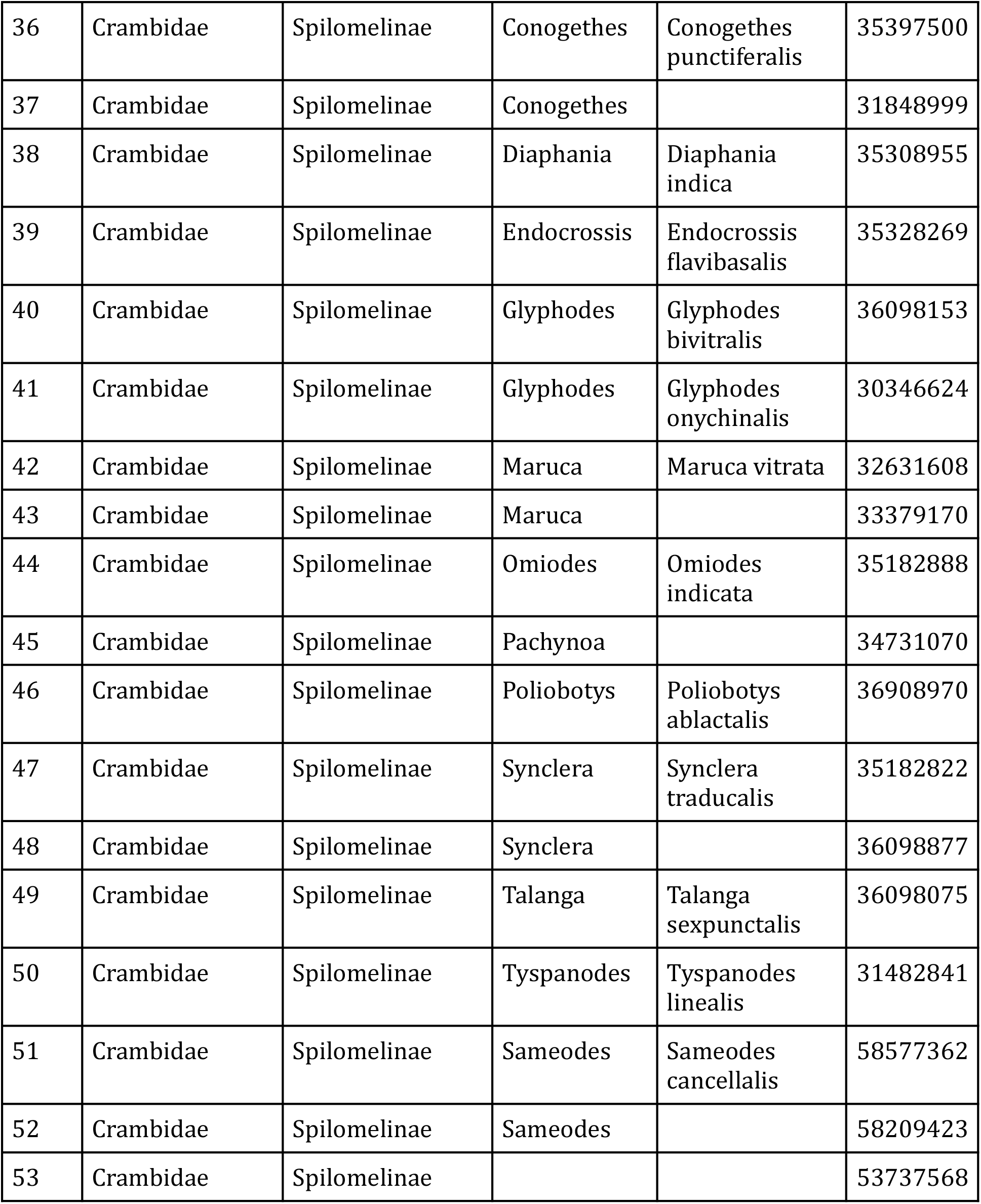
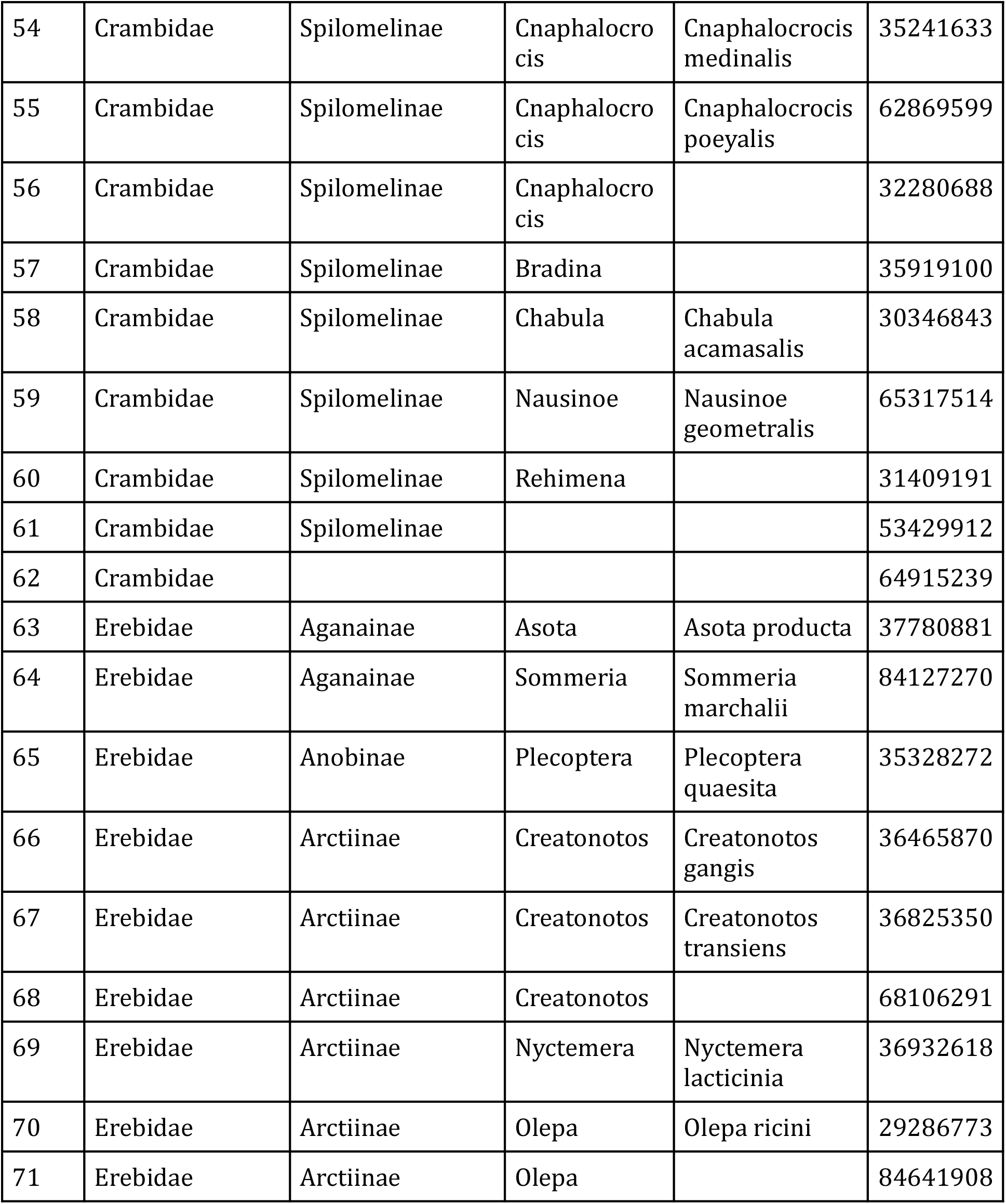
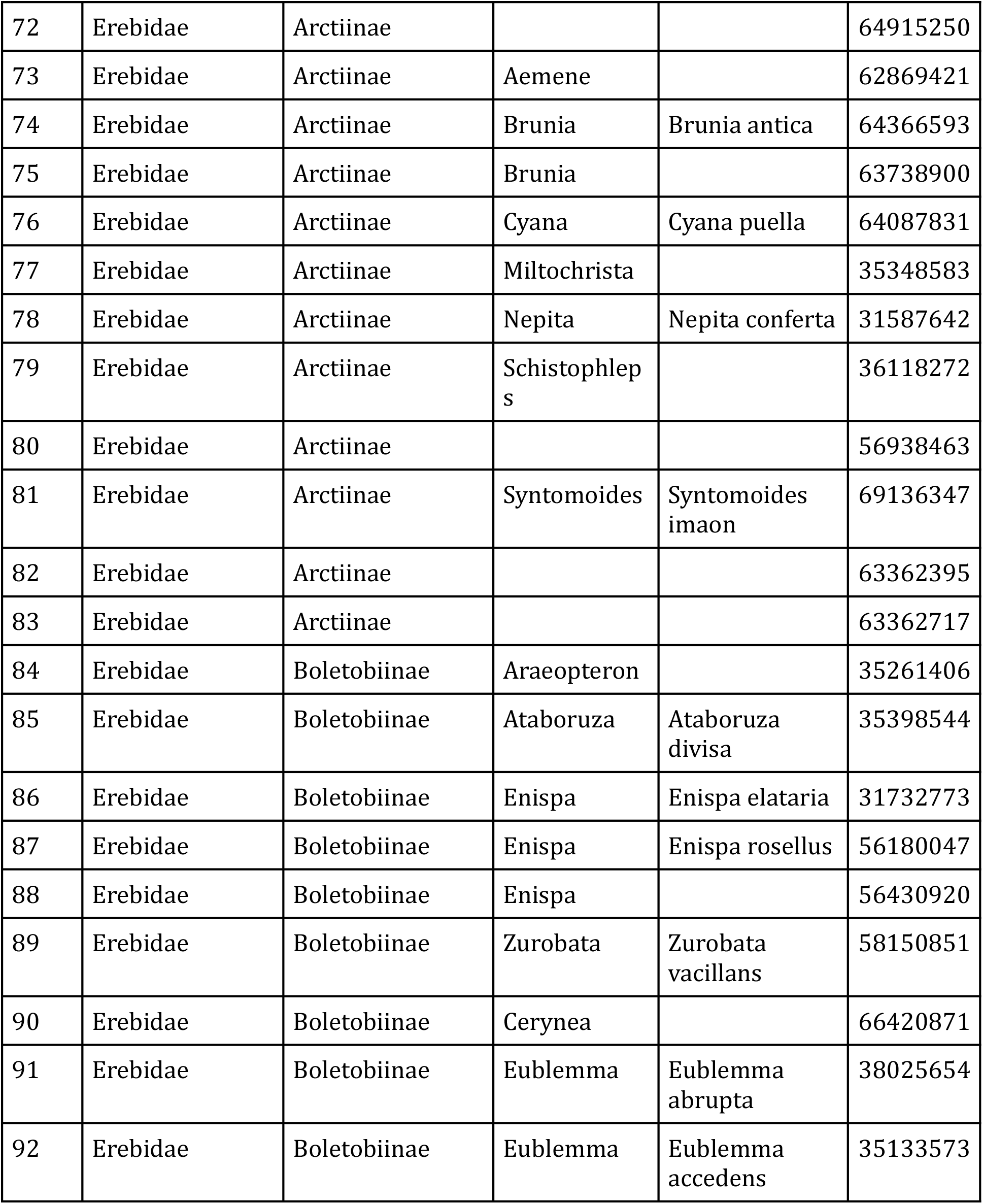
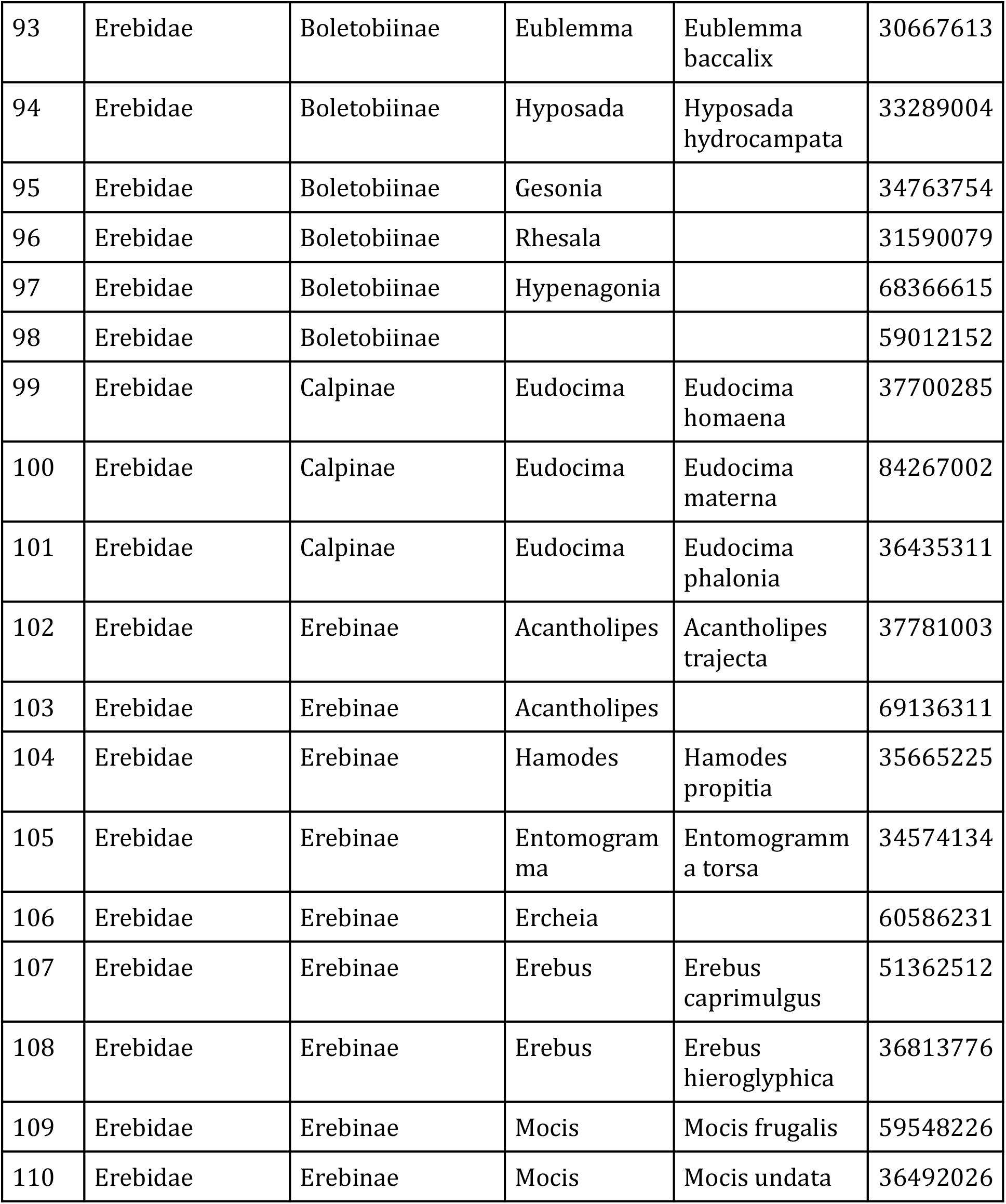
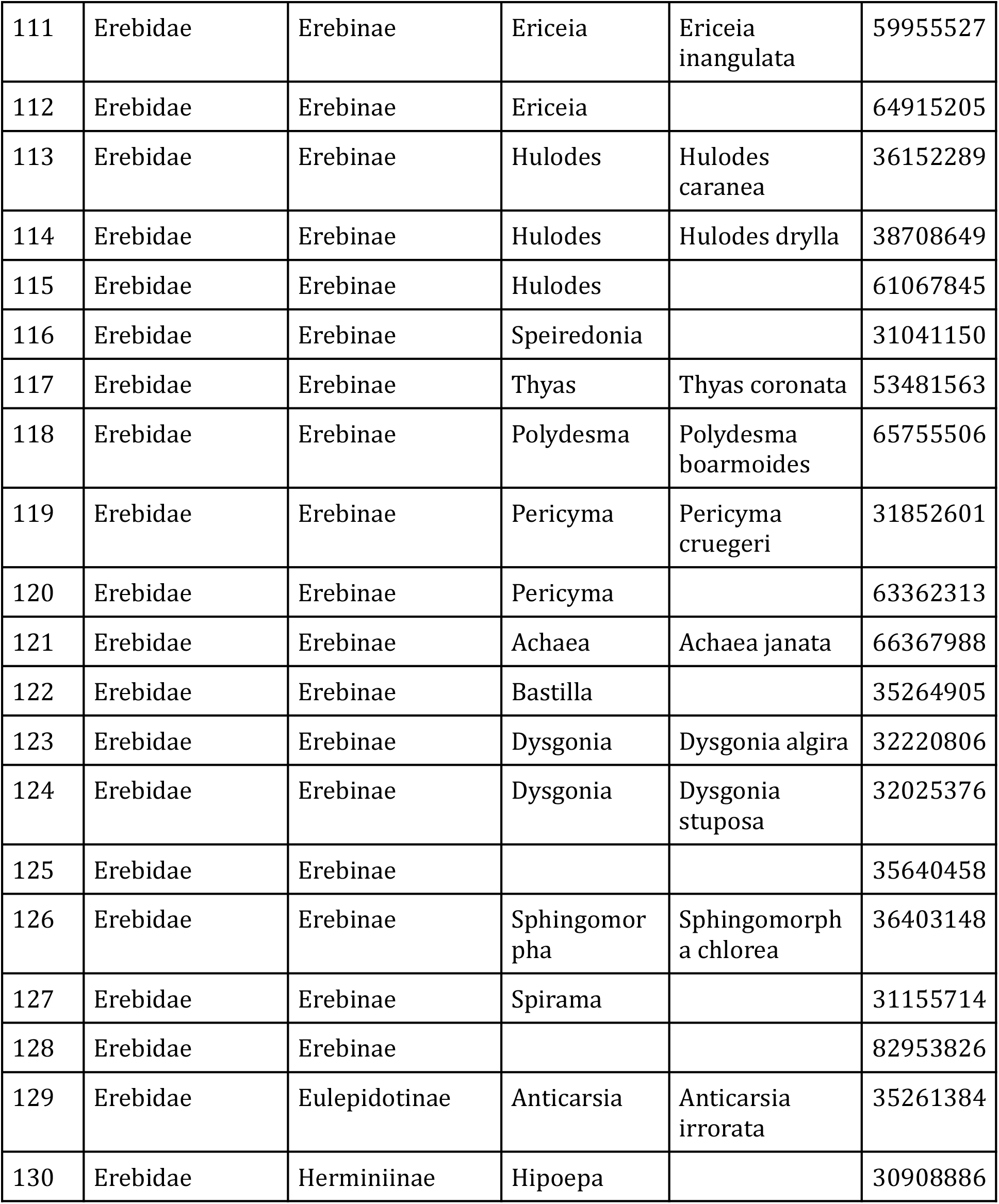
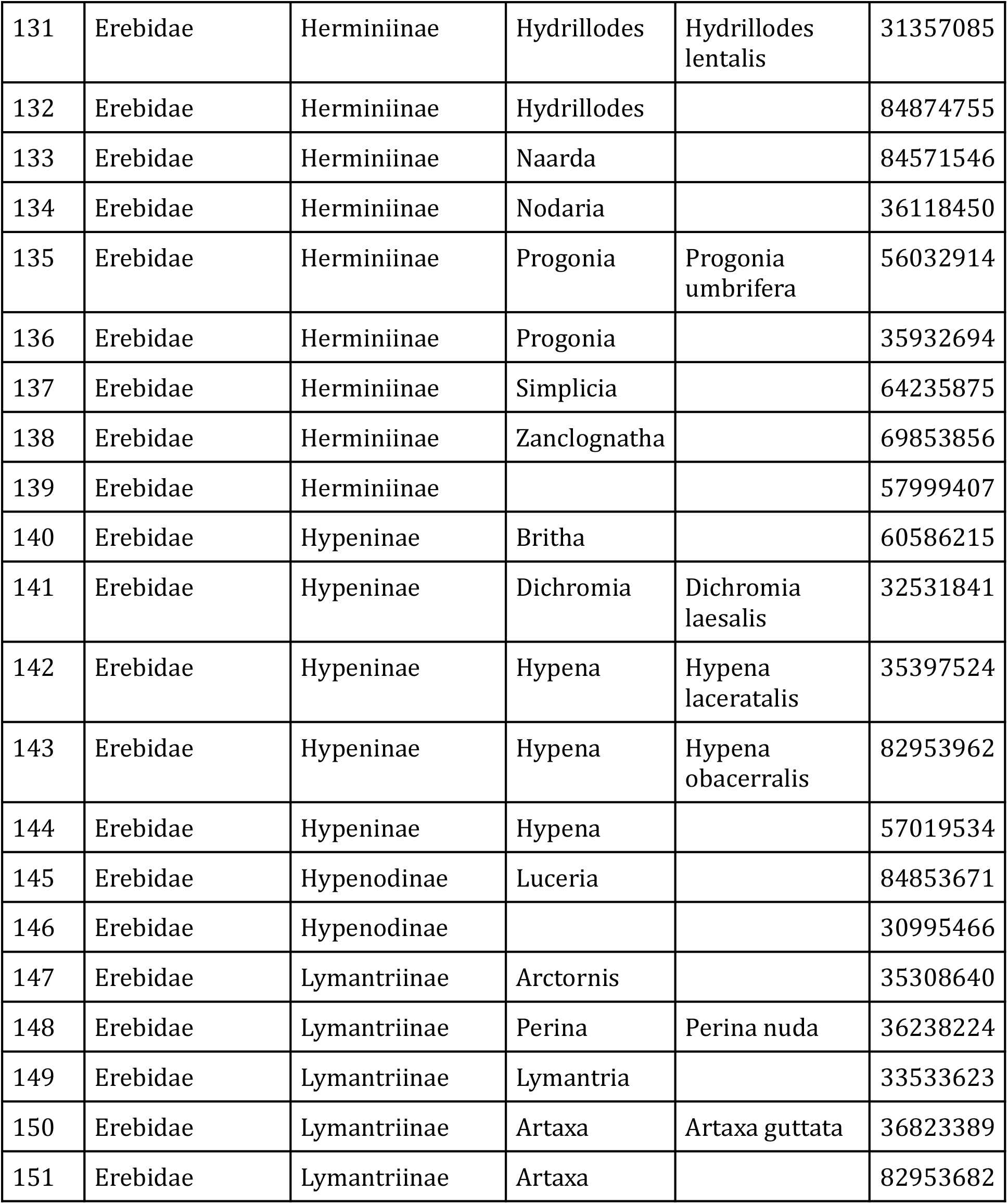
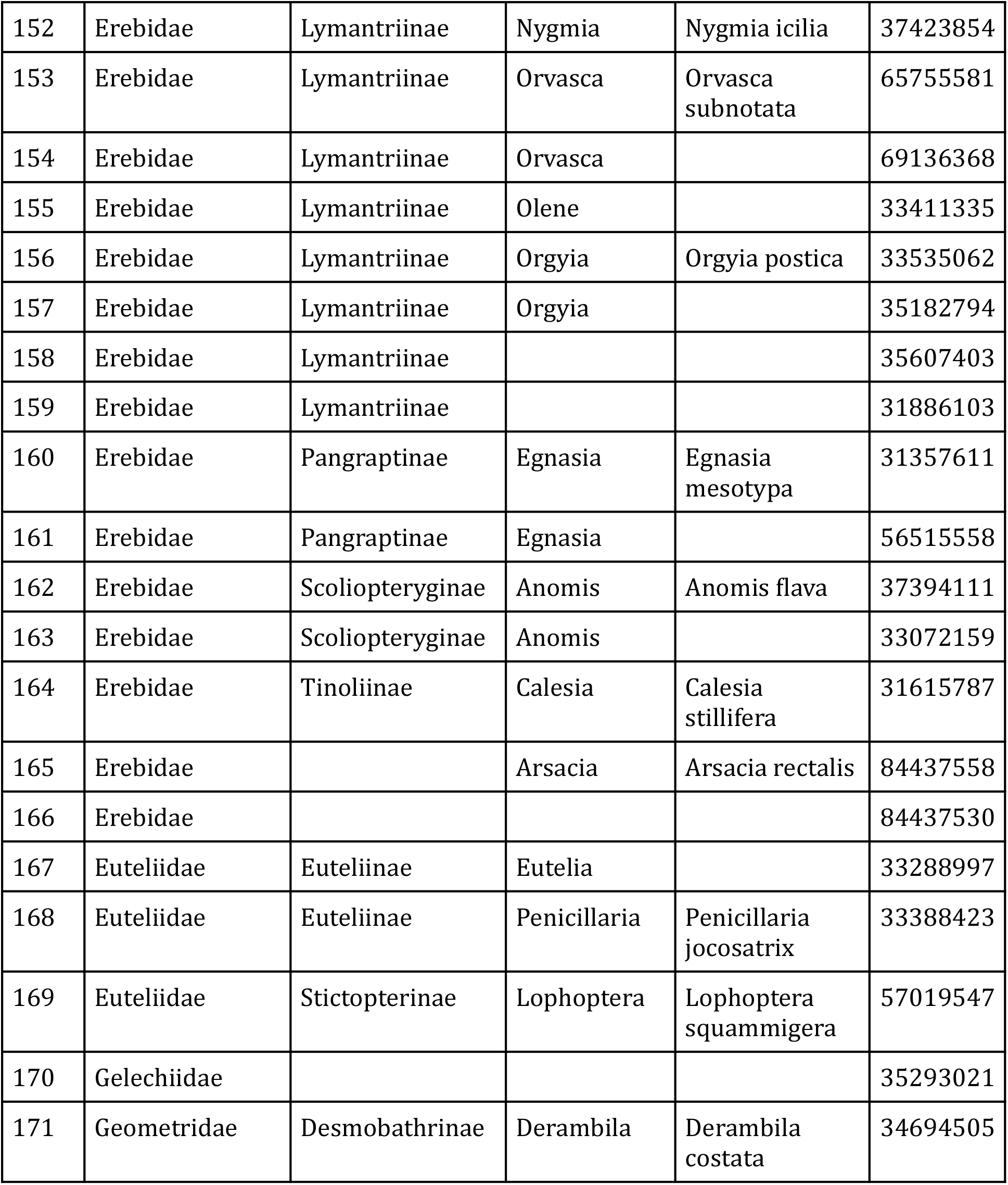
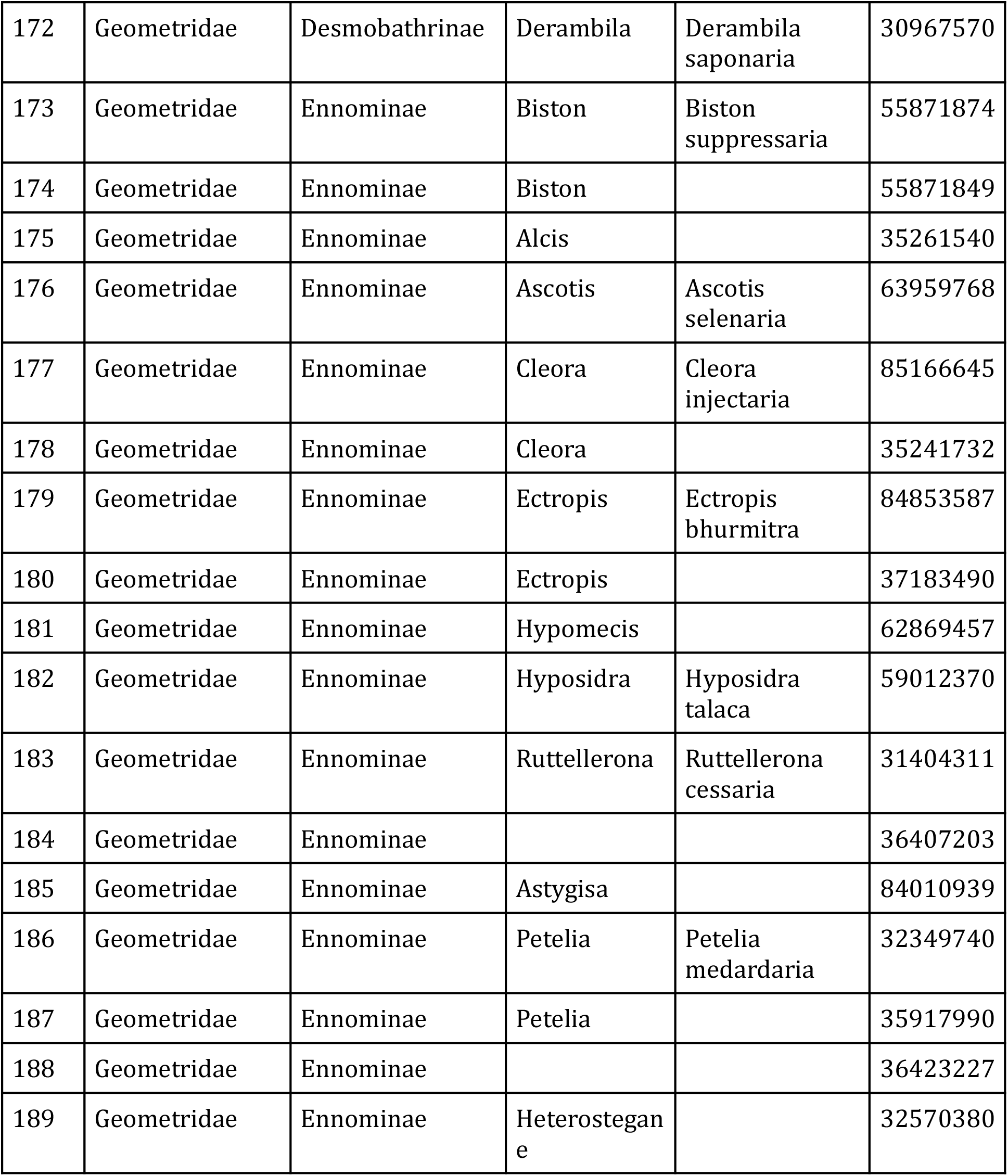
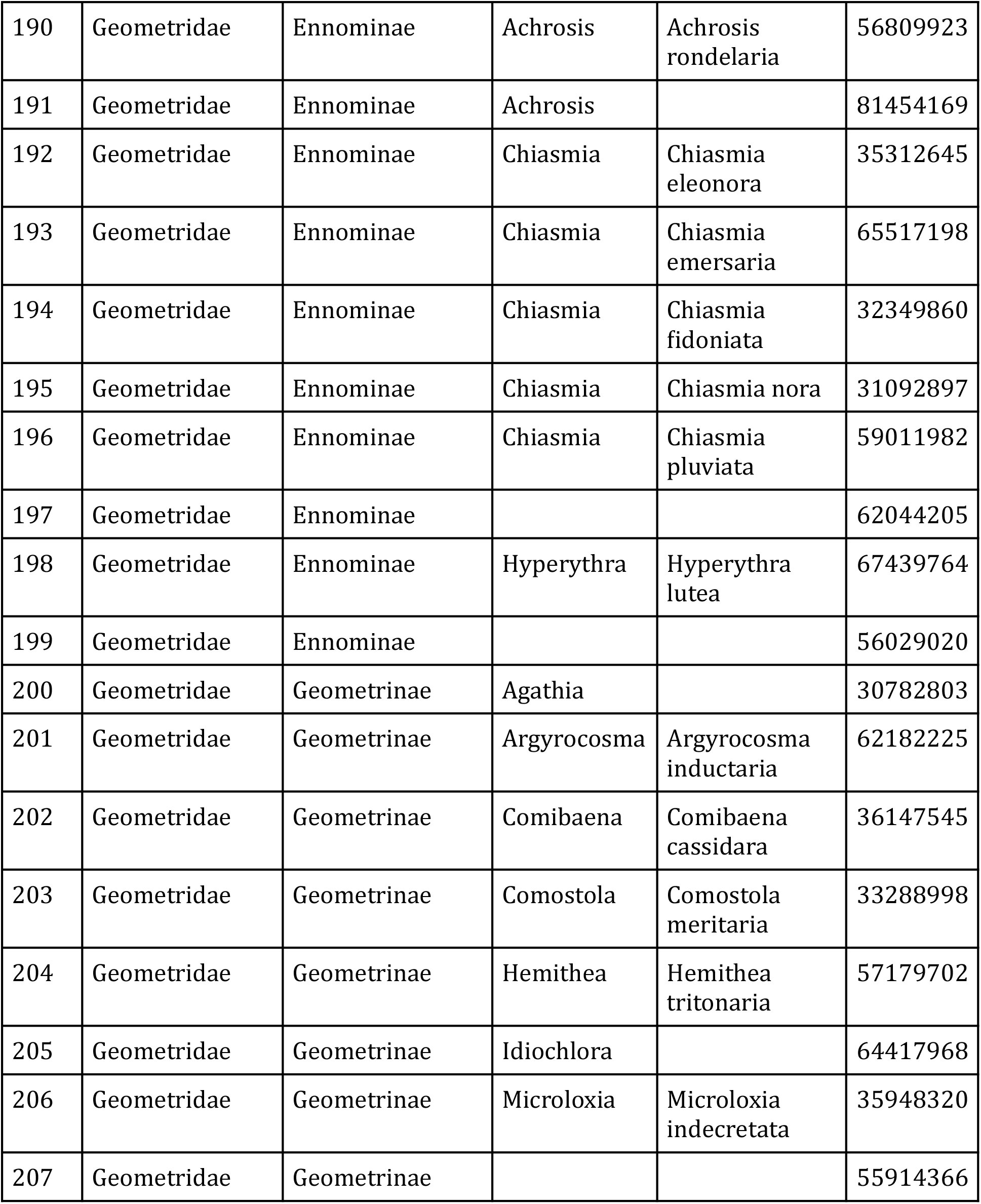
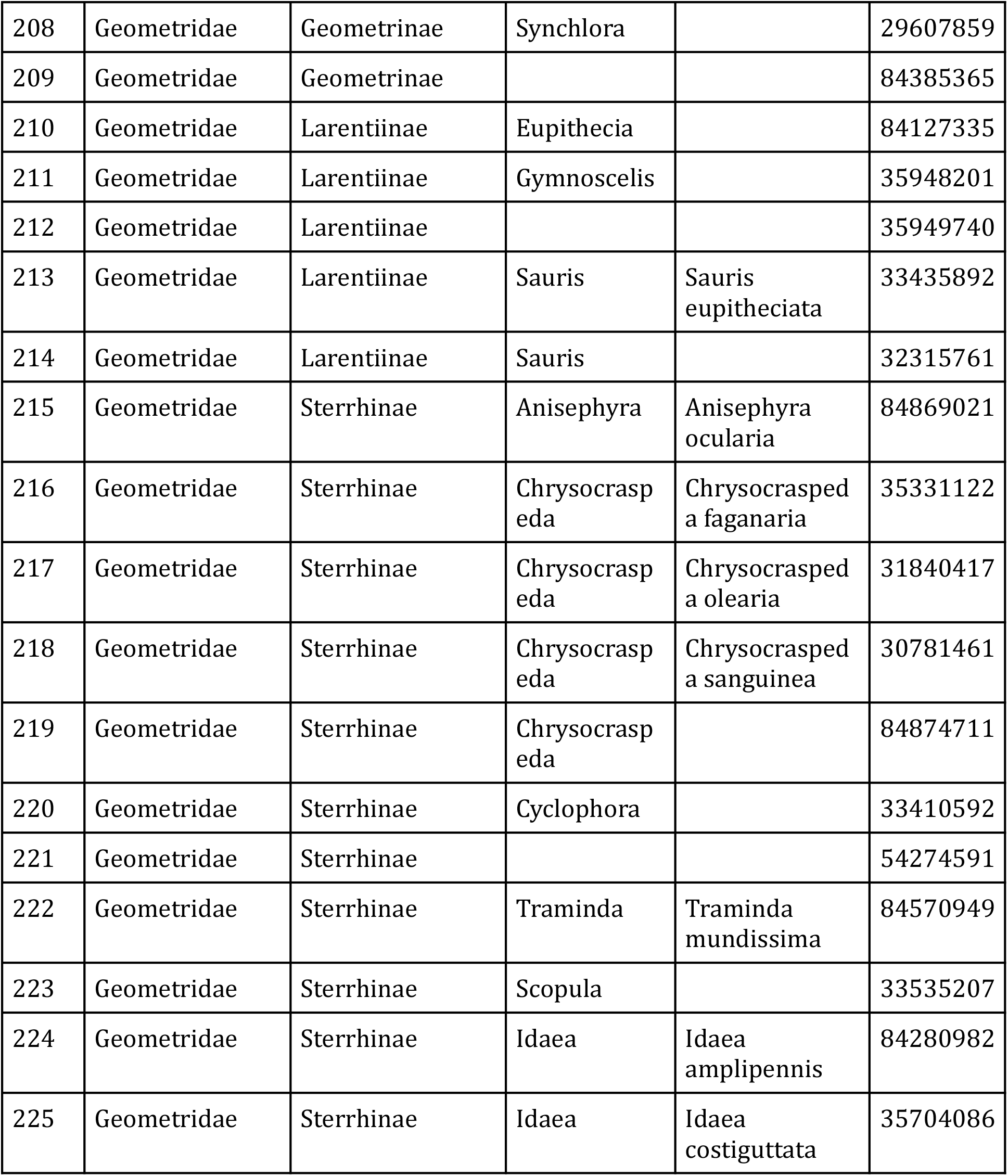
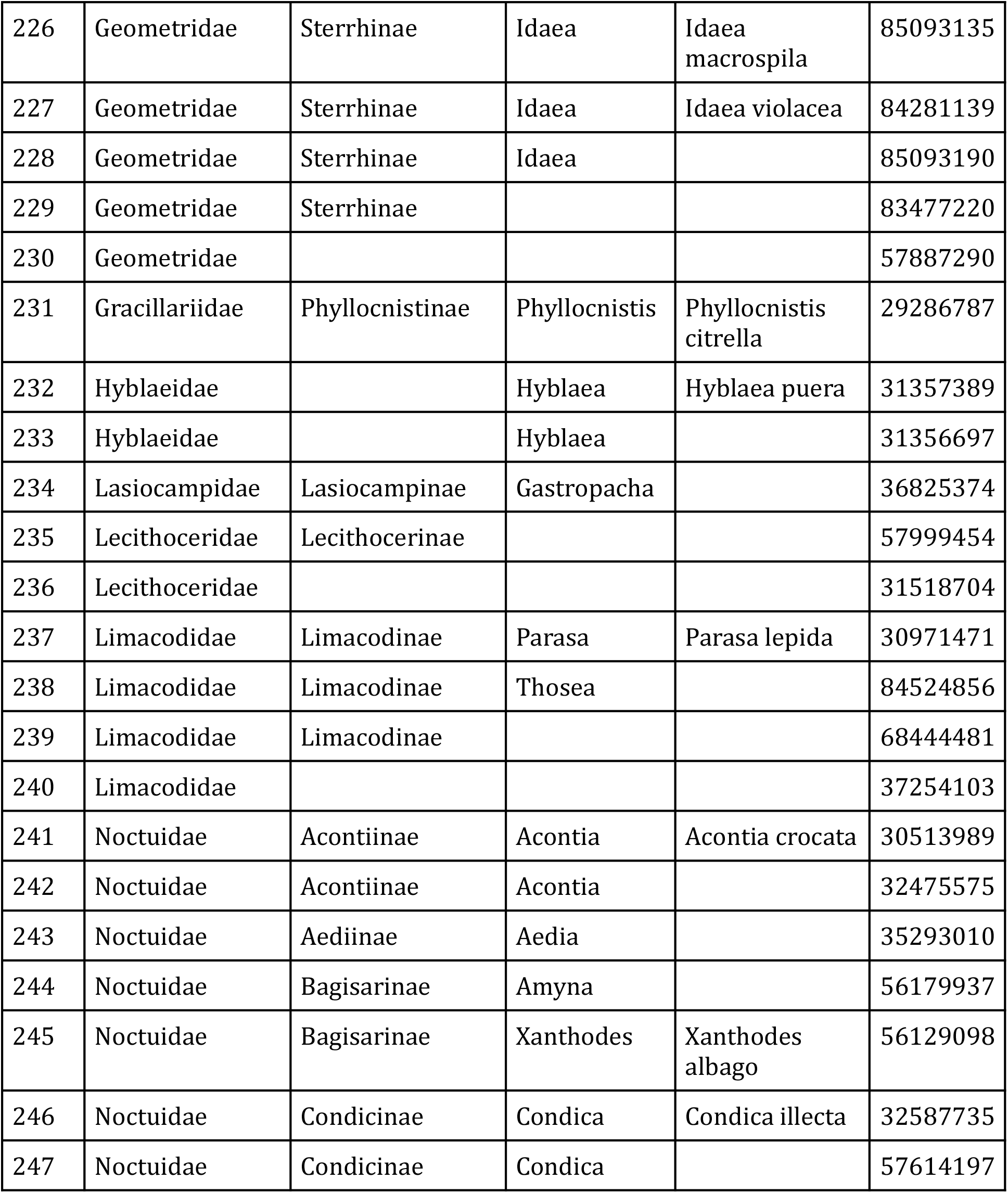
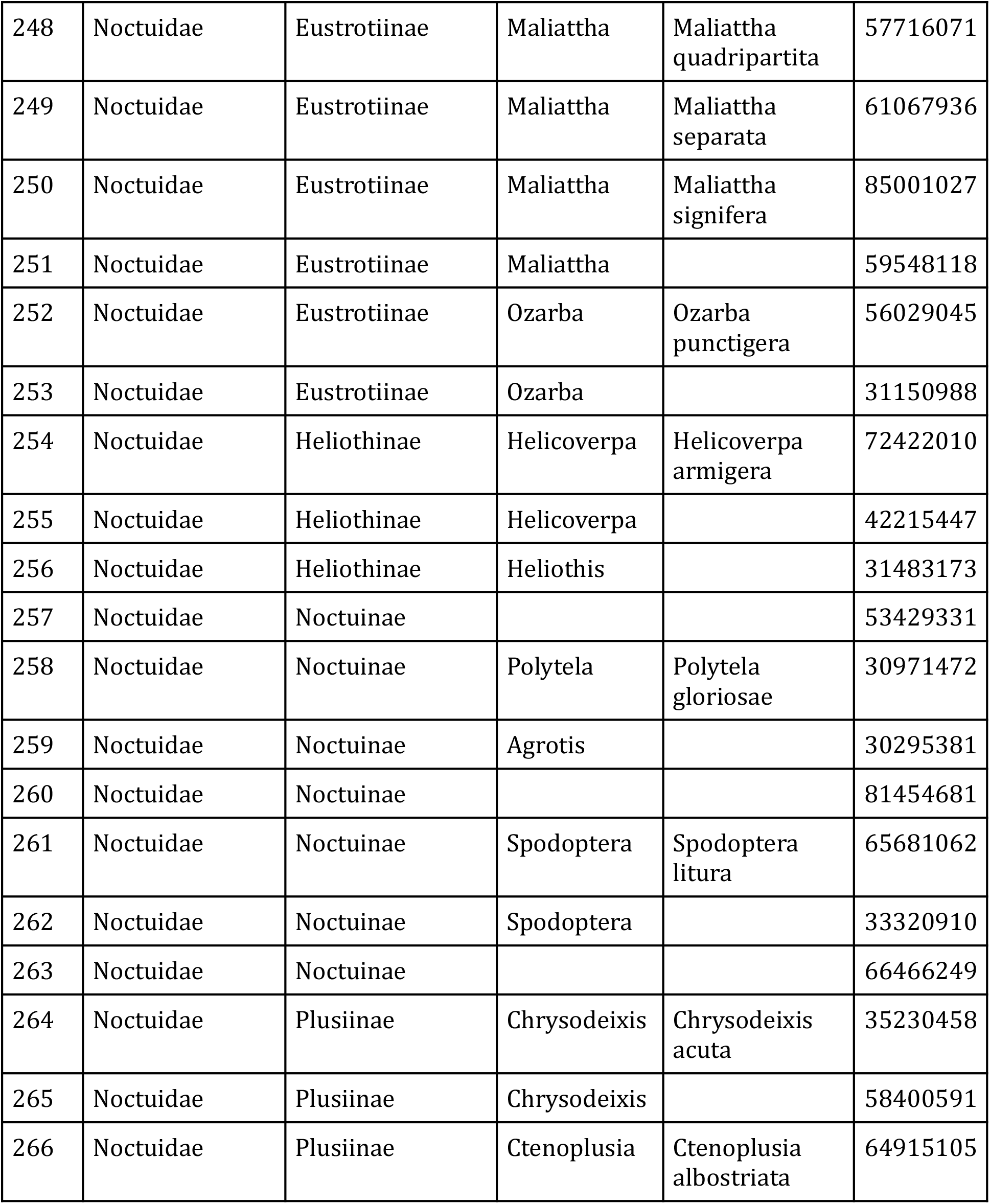
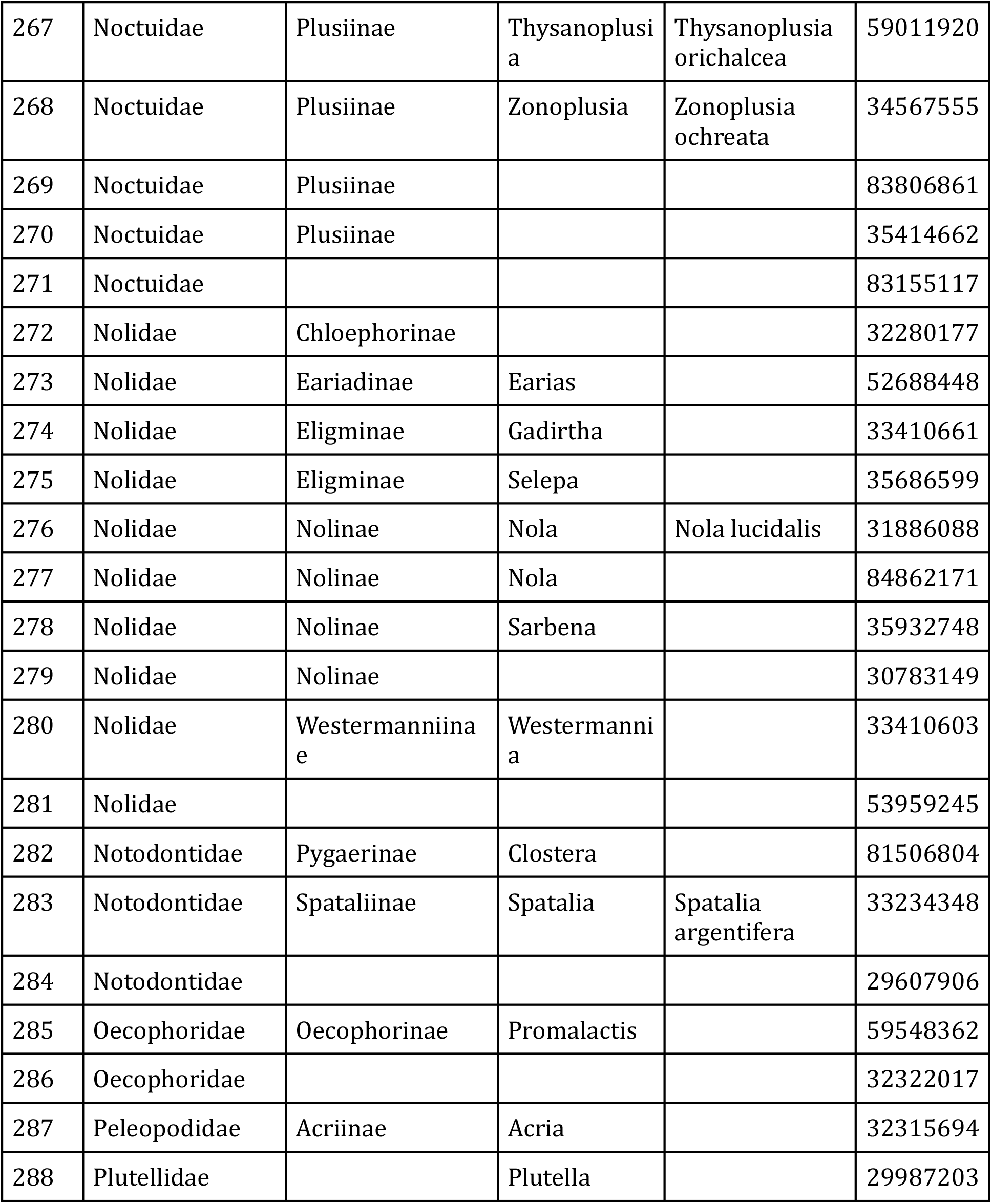
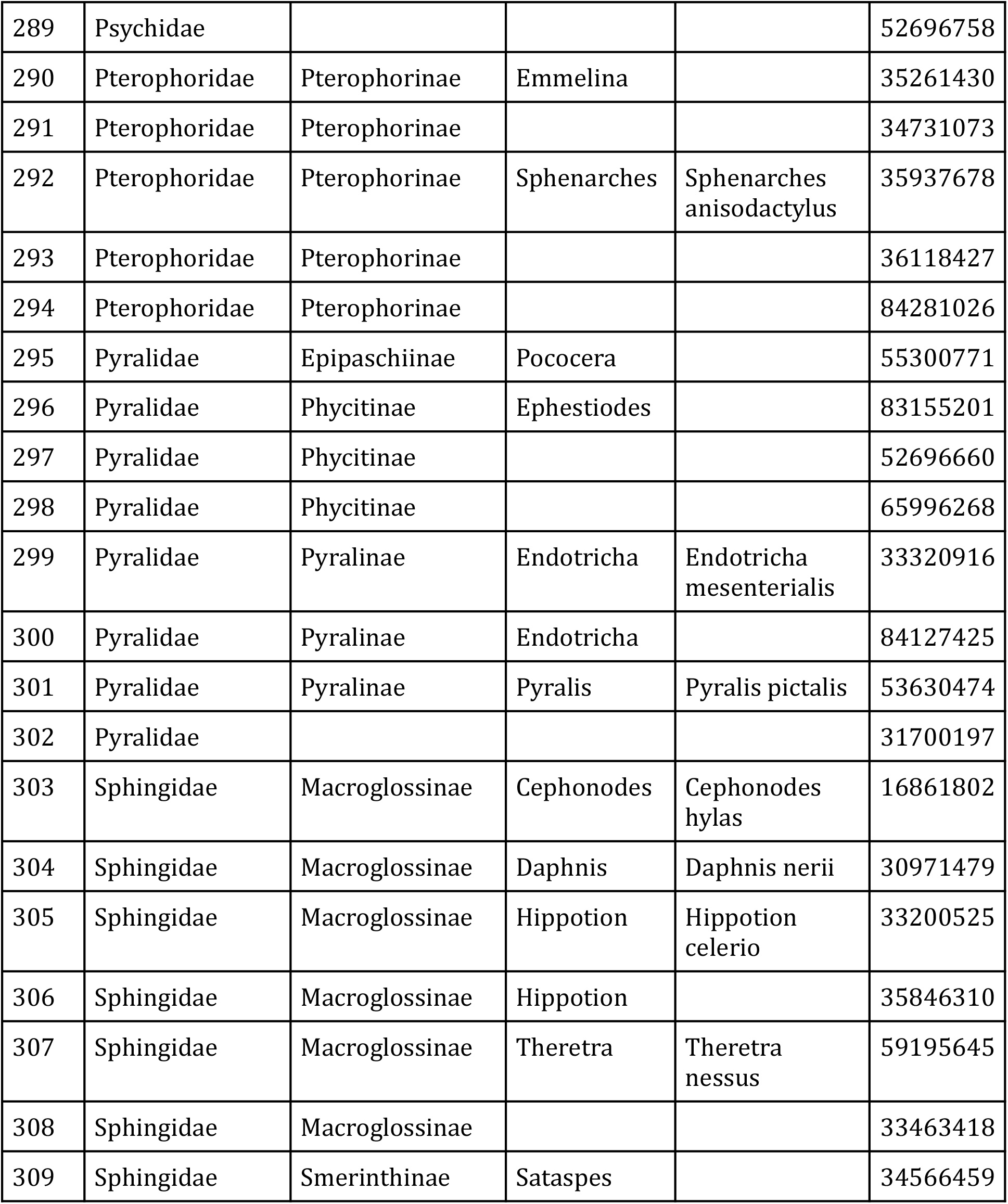
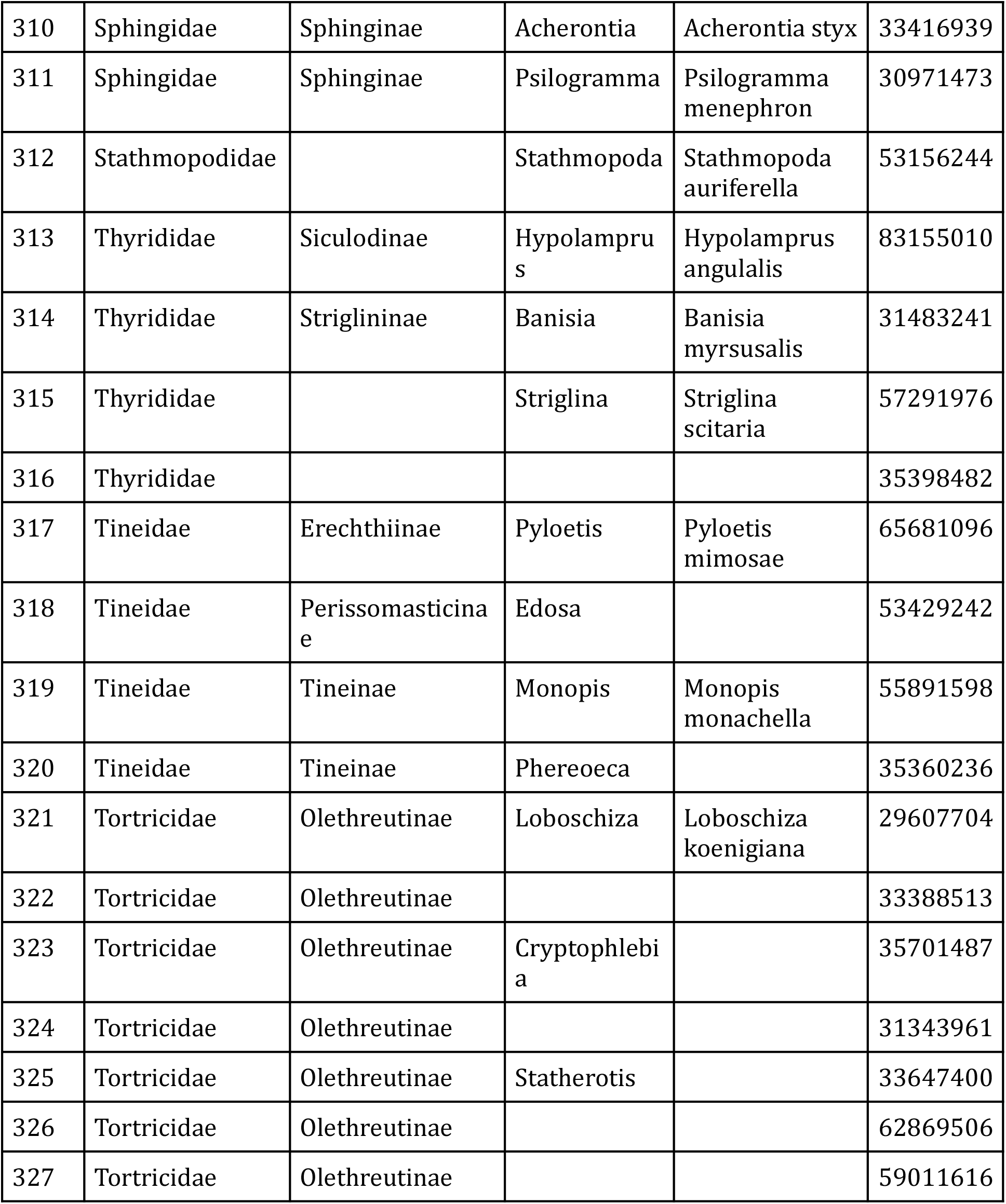
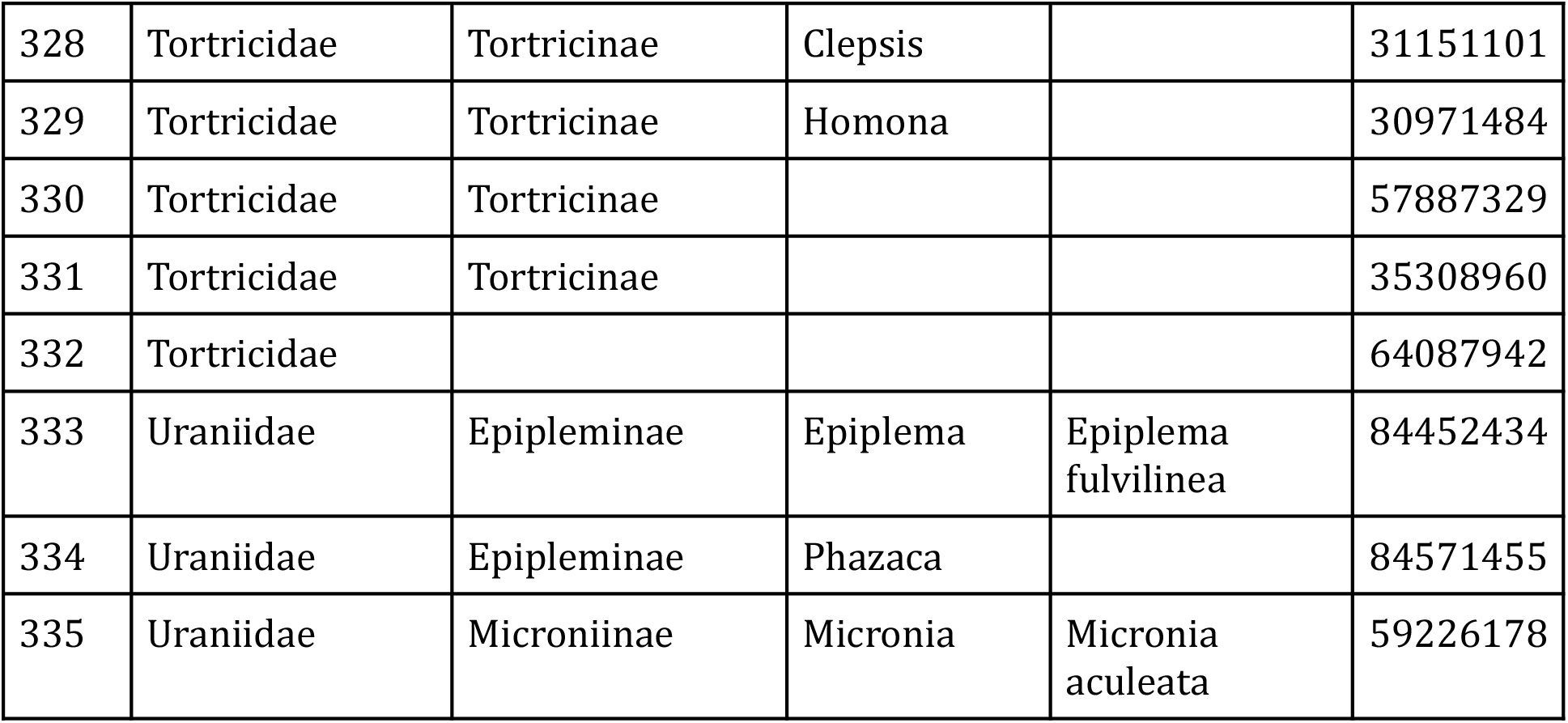
List of moths of Savitribai Phule Pune University, retrieved on Jul 01, 2021.

These observations in total represent 26 families as shown in Fig 2, with five most abundant families being Erebidae (234), Geometridae (147), Crambidae (117), Noctuidae (86) and Pyralidae (23), where the bracketed quantities indicate the observation count. It is worth contrasting the reports to that of Shubhalaxmi et al (Shubhalaxmi (2011)). Both Erebidae, Geometridae families are most abundant in their observations too, however their third abundant family Sphingidae - with 45 species of their 418 total - is considerably absent with merely 13 observations of 7 species.Similarly in the campus of Goa university (Goa, India) only 9 species of Sphingidae have been reported (Gurule and Brookes (2021)). It is thus reasonable to assume that the urban environment does not suit the host plants required for the Sphingidae.

**Fig 2:**
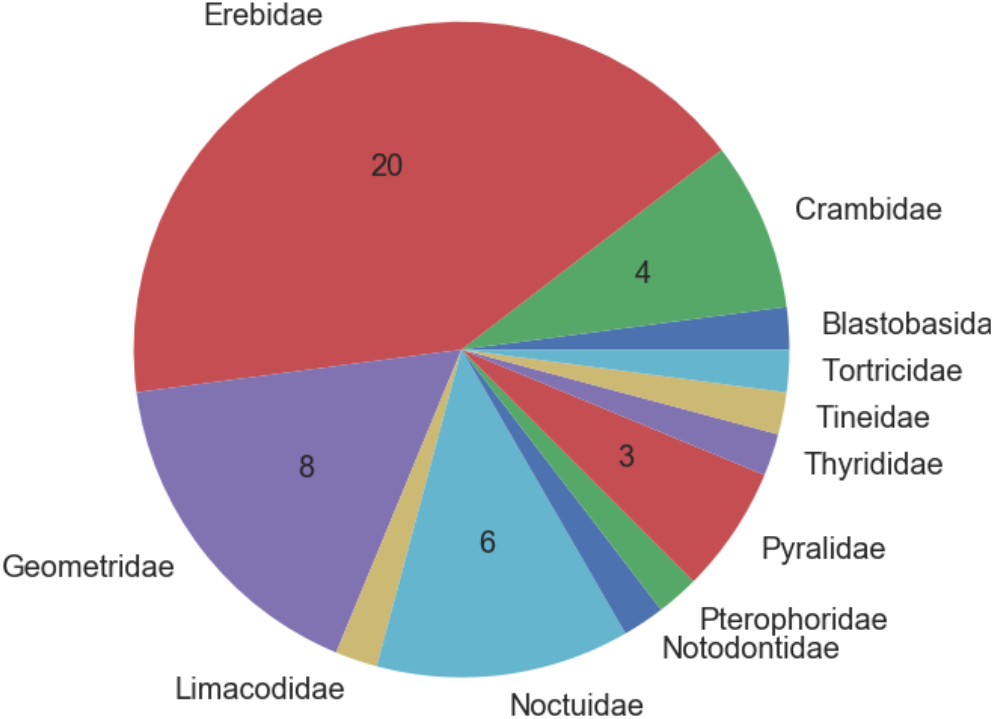
Distribution of families in observed. The numbers indicate uniquely identification of species. Count smaller than 2 are not shown for sake of clarity.

On the other hand, there are some interesting species seen that are rarely reported in Pune or even in India. For example: Spatalia argentifera (Obs ID: 32642126), Sauris eupitheciata (Obs Id: 33435892), Idaea costiguttata (Obs Id: 35704086), Ascotis selenaria (Obs Id: 63959768), Entomogramma torsa (Obs Id: 34574134), Hulodes drylla (Obs Id: 38708649), Hyposada hydrocampata (Obs Id: 33289004). Reports of such species are rare in India and it is indeed hearting to see their presence on the campus. The other notable species is Sommeria marchalii. At the time of preparing this manuscript 49 of 54 observations from India are from Maharashtra state, specifically from the narrow region of western ghats. Pune being in the same region, we have total of eight observations of the species. It appears that this species is fairly localised in west and south of India. Arsacia rectalis is also fairly abundant on the campus. It is reported that the larvae of the species feeds on the plants of Dalbergia genus. It can be speculated - but remains to be verified - that they also feed on Dalbergia melanoxylon which as mentioned above is very common on the campus.

Below in Table 1, we present the detailed checklist. As suggested before, the list is expected to grow, modify over the period of time and thus we associate a date with it. What is being released here should be considered as a snapshot at the time of release of this document, and for all future purposes readers are encouraged to check the URLs given before.

## Discussion

Despite its presence in a busy metropolis the campus of Savitribai Phule Pune University is home to rich Lepidoptera diversity. With a project spanning for more two years we have compiled a checklist of moths and found the presence of 189 genera and 154 species from 26 families. In an attempt to be precise the identification was crowd sourced via <u>inaturalist.org</u> where each identification is effectively peer-reviewed by global community. The identification automatically and periodically retrieved via a Python program - which is also being released with this manuscript. An advantage of such a system is that any addition of new species, or correction to either identification or taxonomic records get automatically incorporated in future versions of checklist. As a result the checklist that is being presented here is dynamic and subsequent changes can be followed up at GitLab^4^ or [author’s webpge](http://cms.unipune.ac.in/~bspujari/Pune_university_Lepidoptera/)^5^. With the given python program anyone should be able to generate similar checklist for area of their interest.

## Acknowledgments

Being a crowd sourced work the author is grateful to all individuals who have help identify the species. In particular author is grateful to Swanand Kesari, Nagabhushan Jyothi(@nagabhushanjyothi), Vijay Barve, Unnikrishanan (@unnikrishanan_mp), Roger Kendrick (@hkmoths), Rohit MG (@rohitmg), Mangesh Kulkarni and Deepashri Saraf. Special thanks are to @po-po-pro for identification as well as critique of the work, and last but most certainly not the least my huge gratitudes are to Piero Toni (@pierotoni10) for all the help and encouragement.

## Footnotes

2 http://cms.unipune.ac.in/~bspujari/SPPU_Lepidoptera

3 GNU Afferro General Public License (GPL) V-3

4 https://gitlab.com/bspujari/sppu_moths

5 http://cms.unipune.ac.in/~bspujari/SPPU_Lepidoptera

